# Migrant semipalmated sandpipers (Calidris pusilla) have over four decades steadily shifted towards safer stopover locations

**DOI:** 10.1101/741413

**Authors:** David D. Hope, David B. Lank, Paul A. Smith, Julie Paquet, Ronald C. Ydenberg

## Abstract

Peregrine falcons (*Falco peregrinus*) have undergone a steady hemisphere-wide recovery since the ban on DDT in 1973, resulting in an ongoing increase in the level of danger posed for migrant birds, such as Arctic-breeding sandpipers. We anticipate that in response migrant semipalmated sandpipers (*Calidris pusilla*) have adjusted migratory behaviour, including a shift in stopover site usage towards locations offering greater safety from falcon predation.

We assessed semipalmated sandpiper stopover usage within the Atlantic Canada Shorebird Survey dataset. Based on 3,030 surveys (totalling ∼32M birds) made during southward migration, 1974 - 2017, at 198 stopover locations, we assessed the spatial distribution of site usage in each year (with a ‘priority matching distribution’ index, PMD) in relation to the size (intertidal area) and safety (proportion of a site’s intertidal area further than 150m of the shoreline) of each location. The PMD index value is > 1 when usage is concentrated at dangerous locations, 1.0 when usage matches location size, and < 1 when usage is concentrated at safer locations.

A large majority of migrants were found at the safest sites in all years, however our analysis of the PMD demonstrated that the fraction using safer sites increased over time. In 1974, 80% of birds were found at the safest 20% of the sites, while in 2017, this had increased to 97%. A sensitivity analysis shows that the shift was made specifically towards safer (and not just larger) sites. The shift as measured by a PMD index decline cannot be accounted for by possible biases inherent in the data set. We conclude that the data support the prediction that increasing predator danger has induced a shift by southbound migrant semipalmated sandpipers to safer sites.

## 1 INTRODUCTION

Predators affect prey populations not only by ‘direct killing’ (also termed ‘lethal’, ‘consumptive’, ‘density-mediated’ or ‘mortality’ effects; Christianson and Creel, 2014), but by inducing prey to adjust behaviour, physiology, morphology and life history to mitigate the danger (Moll et al., 2017). The adjustments are made behaviourally or by adaptive plasticity (e.g. Domenici et al., 2008), and through natural selection (e.g. Reznick et al., 1990). These ‘non-lethal’ (also termed ‘non-consumptive’, ‘trait-mediated’, ‘intimidation’ or ‘fear’) effects on prey populations can be as strong or even stronger than the effects of direct killing, and together they readily propagate to have further effects on other trophic levels (Terborgh and Estes, 2010; Ohgushi et al., 2012).

The effects of predators on prey populations are beginning to be examined at large scales in nature. Madin et al. (2016) and Myers and Worm (2003) consider the effects on marine fish populations stemming from the great reductions in the abundance of top predators, while Heithaus et al. (2008) document many novel repercussions of changing marine predator abundances (see also Estes et al., 2011; Babcock et al., 2010). Still, much of what is known about these relationships between predators and prey relates to lethal effects (i.e. mortality inflicted directly by predators). In contrast, the non-lethal effects (e.g. influences of predator presence on prey behaviour and morphology; see Madin et al., 2016, Box 1) are not as well studied, despite many advances in the past two decades from the experimental literature (reviewed by Long and Hay, 2012).

In contrast to the ongoing reductions in marine systems, top predators are currently increasing in abundance in some terrestrial systems. In the case of the peregrine falcon (*Falco peregrinus*), migrating, wintering, and breeding numbers all show substantial, ongoing increases that began after the 1973 ban on DDT. The historical population in North America is at estimated at 10,600 − 12,000 breeding pairs, the majority of which (∼75%) bred north of 55° (including Greenland; Cade and Burnham, 2003, p. 6) and migrated to lower temperate and tropical latitudes. Migration counts (McCarty and Bildstein, 2005) and mid-winter counts at temperate latitudes (Ydenberg et al., 2017) show strong ongoing increases of 3 − 7 fold that began in the mid- or late 1970s; increases which have led to a current population estimate of over 60,000 falcons across North America (though this is a very rough estimate COSEWIC, 2017).

This population increase includes a large increase in abundance of peregrines breeding at temperate latitudes along continental flyways, partially as a result of programs releasing captive-bred peregrines into the wild. Releases took place throughout the continent, but the biggest programs established breeding peregrines in the Bay of Fundy, and in Delaware and Chesapeake Bays (and environs; Amirault et al., 2004; Gahbauer et al., 2015; Watts et al., 2015). Especially relevant to this paper is the introduction of peregrines into the Bay of Fundy (summarized in Dekker et al., 2011), in which 178 captive-bred birds were released 1982 − 1993 by the Canadian Wildlife Service. The first breeding of peregrines in the region in at least a half-century (and perhaps longer; see Discussion) was recorded in 1989. Subsequently, active nest sites increased to the current level of about 35, as documented in Figure 1. Merlins (*Falco columbarius*) have also become more abundant (Dekker et al., 2011), though without the aid of any reintroduction program. Watts et al. (2015) describe a very similar history in Delaware and Chesapeake Bays. These breeding peregrines are especially significant, for they are present and actively hunting (Dekker et al., 2011) throughout the sandpiper passage period, whereas migratory peregrines do not arrive until late September just as semipalmated sandpiper passage is ending (see Figure 5 in Lank et al., 2003).

**Figure 1.**
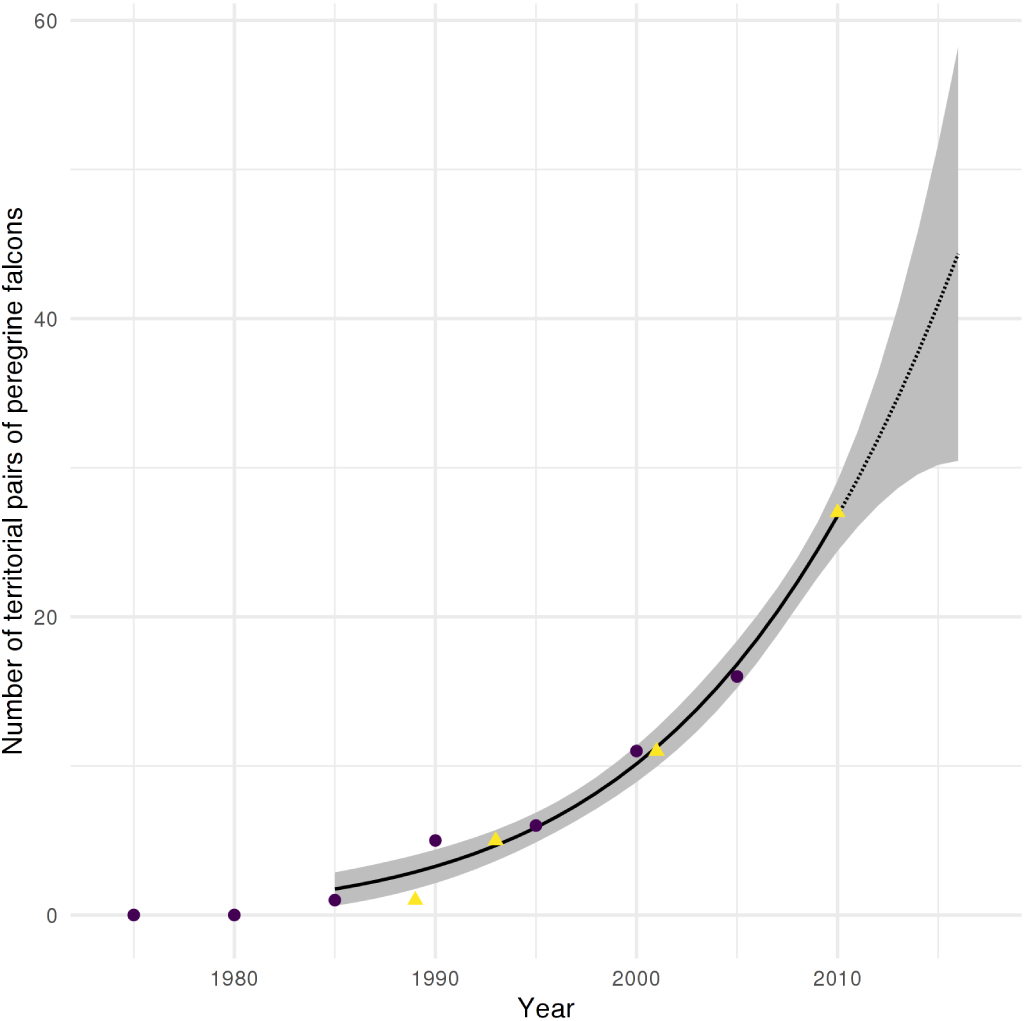
Number of breeding pairs of Peregrine falcons (*Falco peregrinus* in the Bay of Fundy region between 1980 and 2010. Dotted line shows the extension of the quadratic fit curve after 2010. Data from COSEWIC (2017, purple circles) and Dekker et al. (2011, yellow triangles).

**Figure 2.**
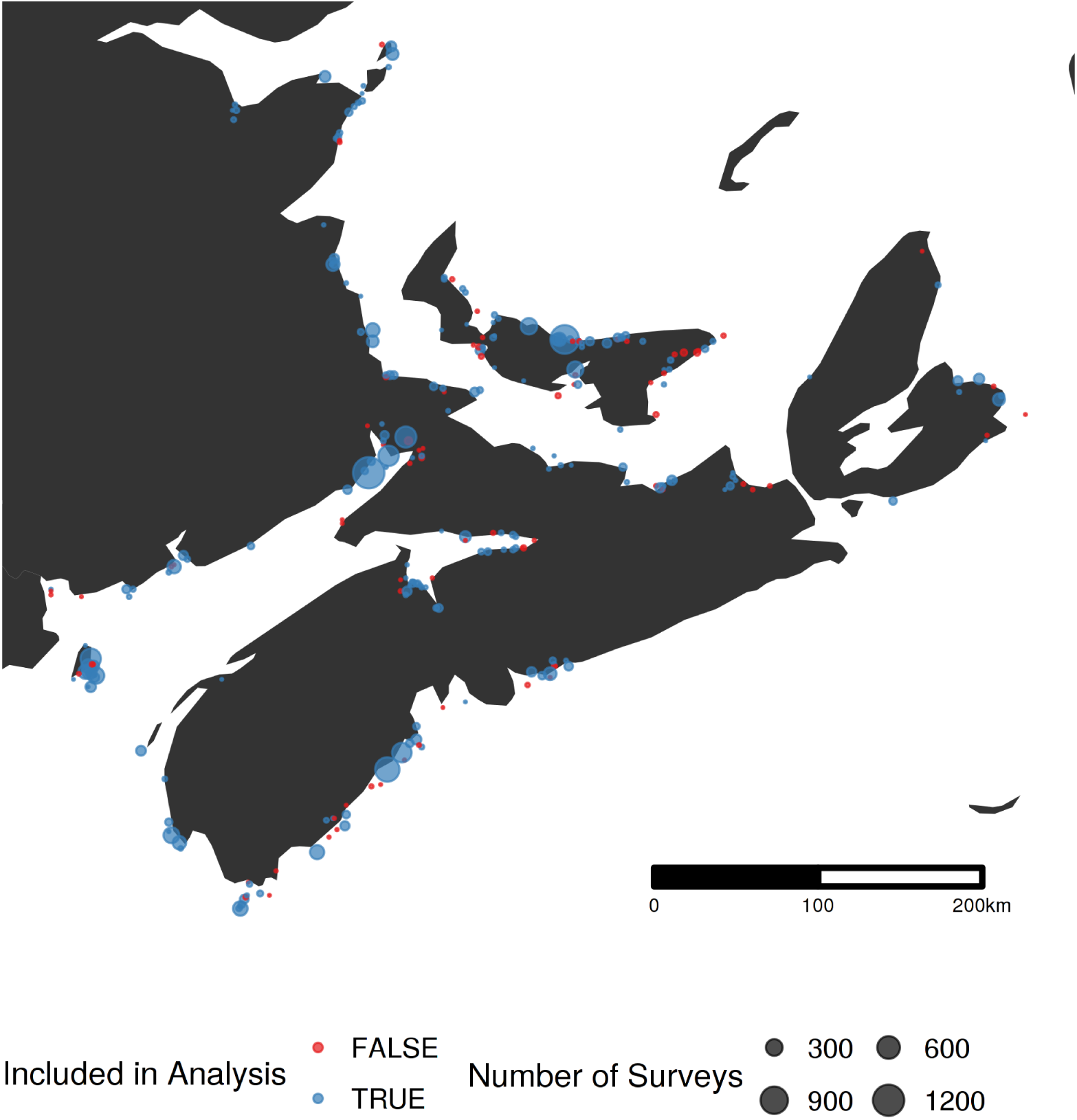
Map of survey locations (n = 198) from the Maritimes portion of the Atlantic Canada Shorebird Survey. The size of each point is related to the number of surveys conducted at that site. Sites excluded from the final analysis are shown in red (n = 77).

**Figure 3.**
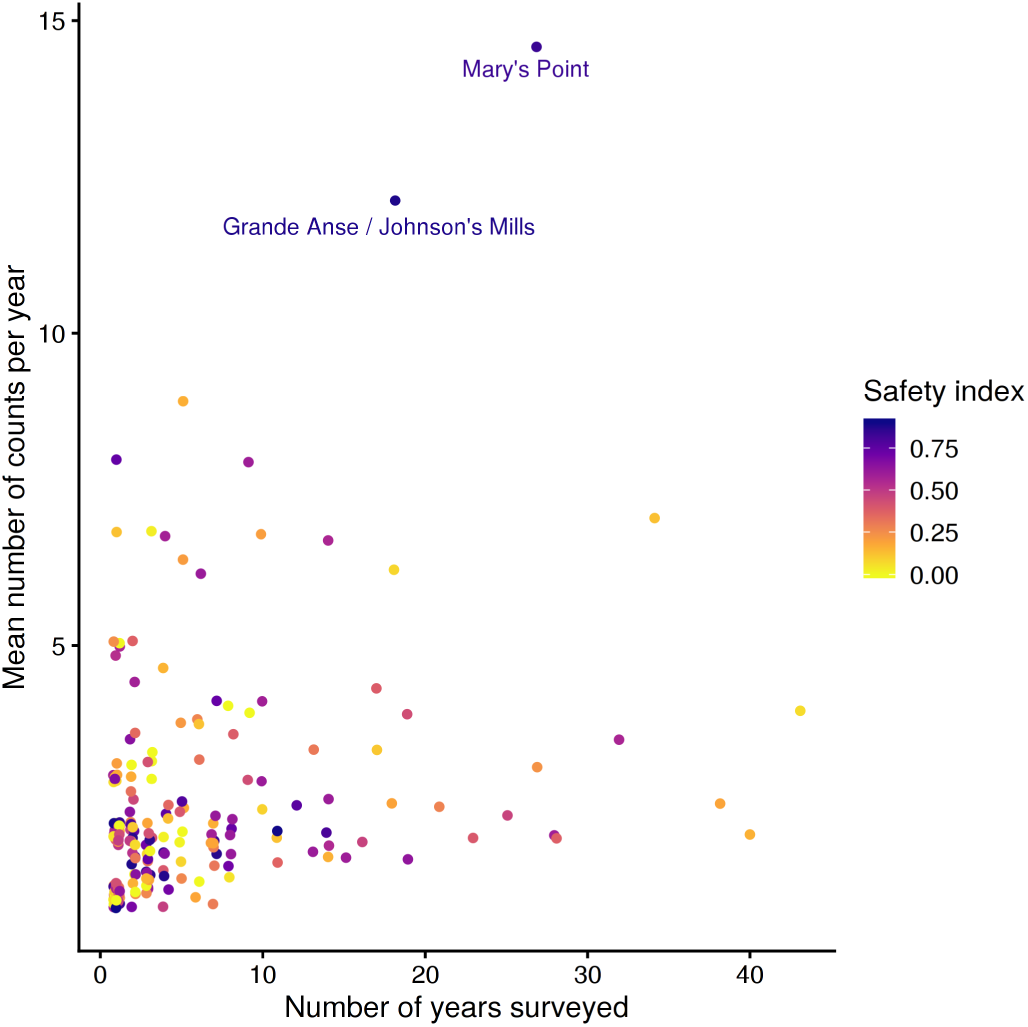
Number of years surveyed and the number of surveys per year for each location. The colour of the point is the safety index (i.e. proportion of the site within 150m of cover; see Methods). The two largest sites are named. Figure S2 shows results of sensitivity analysis around the importance of these two sites.

**Figure 4.**
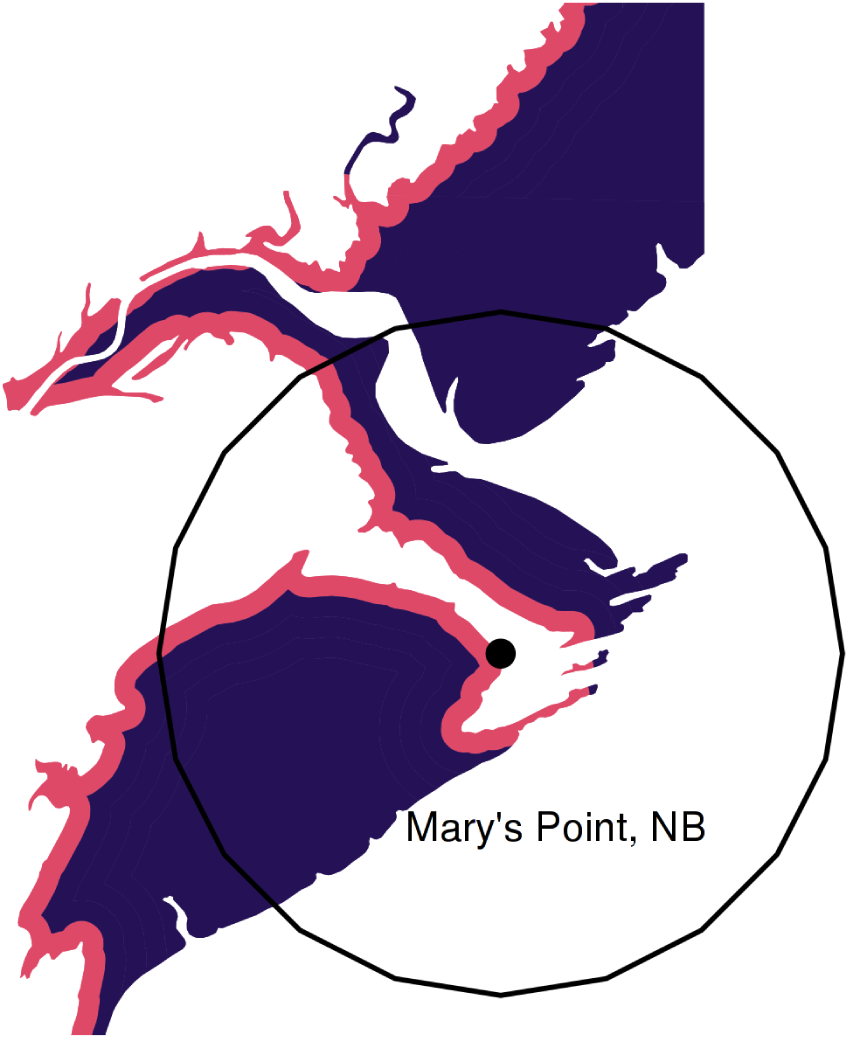
Example showing the safe (purple) and dangerous (red) portions of a habitat. Mary’s Point, NB is shown with its geographic location highlighted by the point. The 2,500 m radius habitat circle is shown around this point. Only intertidal mudflat habitat is shown.

**Figure 5.**
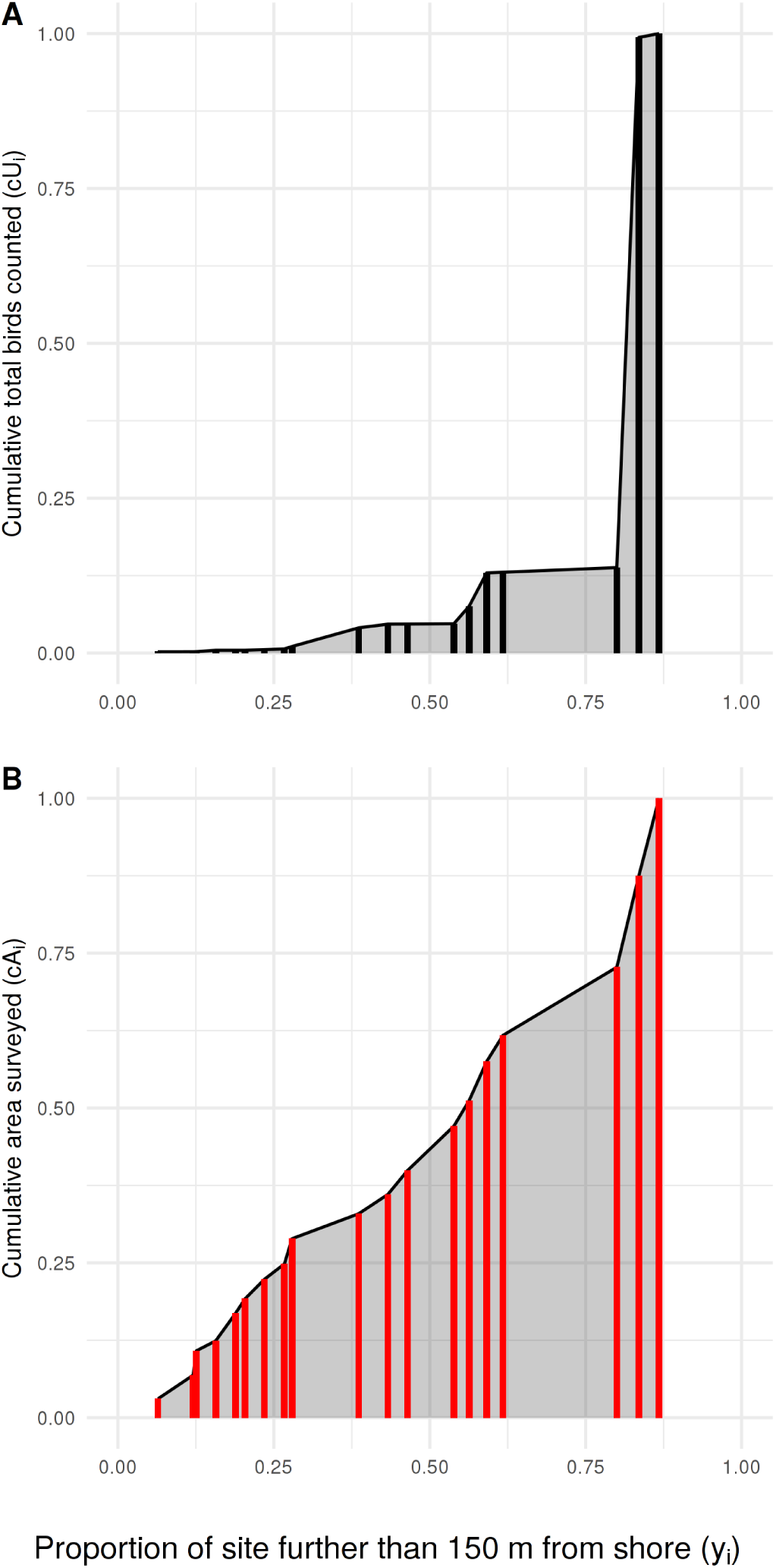
How the Priority Matching Distribution index is calculated, using 1985 in this example. The locations (red and black bars; locations with equivalent safety are stacked) surveyed in a year are ordered along the x-axis from lowest to highest safety index (*y*_*i*_), with the cumulative proportional usage (Panel A: numerator of the PMD index), and intertidal area (Panel B: denominator of the PMD index) shown by the height of the vertical bar. The grey area in each plot shows the area under the distribution (AUD) used to calculate the PMD index.

As a result of these increases, southbound migration along the Atlantic coast of North America has become much more dangerous for sandpipers. We assert that the ongoing recovery of falcons and other raptors constitutes an important environmental change for many prey species that should induce strong risk effects. Demonstrated non-lethal consequences of the increased exposure to raptors include seabirds shifting to safer breeding locations in response to the recovery of bald eagles (*Haliaeetus leucocephalus*; Hipfner et al., 2012). The increased abundance of white-tailed sea eagles (*Haliaeetus alba*) in the Baltic Sea caused barnacle geese (*Branta leucopsis*) to alter migration timing and to shorten the duration of parental care (Jonker et al., 2010). Falcon recovery drove Pacific dunlins (*Calidris alpina pacifica*) to lose their mid-winter fat reserve, and to take up over-ocean flocking in place of roosting at high tide (Ydenberg et al. 2010; see also Dekker et al. 2011]). Of special relevance here is the demonstration that dunlins redistributed during the non-breeding season, shifting towards greater aggregation at safer sites (Ydenberg et al., 2017). In this report we focus on semipalmated sandpipers (*Calidris pusilla*) migrating southward through the Bay of Fundy. We analyze a large dataset of migratory censuses, predicting that stopover site usage has shifted toward greater use of safer sites, analogous to that found for non-breeding dunlins. As demonstrated by Ydenberg et al. (2004; Pomeroy et al., 2006), safer sites are those at which sandpipers can feed distant from shorelines, where cover provides falcons the opportunity for stealth hunts (Dekker and Ydenberg, 2004). Stealth hunts are far more successful than open hunts, which often require peregrines to undertake lengthy pursuits (though see Cresswell and Quinn, 2013, for a contrasting prespective with multiple predators).

Semipalmated sandpipers display many of the attributes expected of mortality-minimizing migrants (Hope et al., 2011; Duijns et al., 2019), and hence we expect that safety is important to their stopover decisions. Small shorebirds show a diverse range of behavioural tactics in response to predation danger (e.g., Lank et al., 2003; Sprague et al., 2008; Pomeroy et al., 2008; Fernández and Lank, 2010; Beauchamp, 2010; Hilton et al., 1999; Martins et al., 2016; Ydenberg et al., 2004; van den Hout et al., 2010, 2017; Cresswell and Quinn, 2013, and references therein). Most of a migrant’s time is spent at stopover sites (Hedenström and Alerstam, 1997), foraging to acquire the fuel necessary for extended flights (Houston, 1998; Cimprich et al., 2005). From the Bay of Fundy, many semipalmated sandpipers make a long (∼4000 km or more) trans-Atlantic flight to South America, acquiring a fuel load nearly equal to their (lean) body mass to do so. The intensive foraging required to accumulate the fuel load compromises vigilance level (Beauchamp, 2014), and a large fuel load makes migrants more vulnerable to predator attack. Characteristics of stopover sites, such as the amount of concealment cover available to predators, or the distance between this cover and the feeding sites used by shorebirds, make the intrinsic danger of some sites higher than others (Lank and Ydenberg, 2003).

There are hundreds of potential stopover sites along the Atlantic coast, and in selecting stopovers shorebirds must balance the risk of predation with the benefits of good foraging conditions. In general, it appears that safety and food trade off at stopover sites so safe sites with high food availability are rare or non-existent. Previous studies have demonstrated that migrating sandpipers avoid sites that do not provide some element of food and safety (Sprague et al., 2008; Pomeroy et al., 2008). An increase in predator abundance is expected to shift the balances of these risks and rewards, leading to a shift away from dangerous stopover sites and towards safer ones (Hope, 2018).

Conditions other than the level of predation danger have also changed over recent decades for migrant sandpipers. These include the degradation of existing and the appearance of new habitats (Iwamura et al., 2013; Studds et al., 2017; Taft and Haig, 2006; Alves et al., 2012), climate change (Both and te Marvelde, 2007; Gordo, 2007; Cox, 2010; Sutherland et al., 2015; Mann et al., 2017), and possible strong population reductions (Munro, 2017; Rosenberg et al., 2019). These changing conditions could also affect stopover usage by alterations in site characteristics, energy requirements, or the degree of competition.

We use a large survey dataset of counts of southbound semipalmated sandpipers at Bay of Fundy stopover sites to determine if semipalmated sandpipers changed site usage between 1974 and 2017. We developed an index to describe the annual distribution of birds and utilized statistical methods and simulations to diagnose whether semipalmated sandpipers adjusted stopover site selection as predator abundance increased. We predict a shift in bird usage towards safer sites as migrants increasingly had to prioritize safety.

## 2 METHODS

### 2.1 Study region

The semipalmated sandpiper is a shorebird species with a hemisphere-spanning migration (Brown et al., 2017). The breeding range stretches across arctic North America, and the non-breeding range across the northern coasts of South America (Hicklin and Gratto-Trevor, 2010). Migrants moving from the breeding grounds pass either through the interior or to the east coast of North America. Atlantic Canada holds the most important staging areas (Hicklin, 1987), with numerous potential stopover sites, especially around the Bay of Fundy (Garrett, 1972; Hicklin and Smith, 1984; Sprague et al., 2008; Quinn and Hamilton, 2012). Migrants arrive from the central and eastern portions of the breeding range, load large amounts of fuel, and depart to the south-east over the Atlantic Ocean (Lank, 1983),making a single flight of over 4000 km directly to South America (Lank, 1979).

### 2.2 Shorebird Surveys

The Atlantic Canada Shorebird Survey (ACSS) is organized by the Canadian Wildlife Service and has been conducted annually since 1974 to identify important stopover sites for migrating shorebirds and to help assess population trends. Surveyors attempt to census sites every second weekend during the southward migration period. Count methodology is described in detail by Morrison et al.(1994; Gratto-Trevor, Smith, Morrison, Aubry, and Cotter 2012; see also the ACSS survey protocol and guidelines - Environment and Climate Change Canada 2014). Protocols aim to make procedures consistent within sites across years, but there is substantial variability in methodology and effort among sites.

The data for the analysis reported here were accessed through Bird Studies Canada’s Nature Counts database (Environment and Climate Change Canada, 2008). We focused on sites where semipalmated sandpiper were censused 1974-2017 throughout Nova Scotia, New Brunswick, and Prince Edward Island (Figure 2). Sites in Newfoundland (due to their position ancillary to the main semipalmated sandpiper migration route) and those at which semipalmated sandpipers have never been recorded were excluded. We included surveys during the main migratory period, defined as falling within the 10th (July 28) and 90th (August 21) quantiles of all semipalmated sandpipers counted between July and October. After this filtering, our analysis incorporated 3,030 of the 20,064 surveys, and 471 of the 769 survey sites in the full dataset.

Each survey site is associated with a name describing the geographic locality, and a latitude and longitude in decimal degrees. To reduce possible pseudoreplication due to spatial autocorrelation, we pooled sites that were within 375m of each other, reducing the 448 ‘sites’ into 198 ‘locations’. The number of years that each site was surveyed and the number of surveys per year varied widely Figure 3). We used the mean of all the site-surveys in a year at a location, whatever the methodology, to represent that location in order to reduce any bias possibly arising with variation in the number of counts per location.

### 2.3 Site Characteristics

We assigned measures of ‘size’ and ‘safety’ to each location. We used the area of intertidal habitat in a 2500m radius around each location’s geographic point as the measure of location size (Figure 4). For many locations the measure of size is unaffected by the radius, but on large areas, particularly those along a large straight coastal stretch, the amount of intertidal area is strongly dependent on the radius. We chose 2500m based on our own experiences, in which we observed that foraging sandpipers quickly reacted to disturbances occurring within a few kilometres. Foraging sandpipers can traverse this distance in a few minutes (Reurink et al., 2016). We examine the sensitivity of the results to this assumption in Figure S2.

As defined by Lank and Ydenberg (2003) ‘danger’ represents the inherent riskiness of a location (see also Hugie and Dill, 1994). We indexed safety as the proportion of the intertidal area at each location lying beyond 150m of the shoreline, where foraging is most risky (Equivalently, danger is the proportion of a location’s intertidal area lying within 150 m; Pomeroy, 2006; Dekker and Ydenberg, 2004; Pomeroy et al., 2008). The 150m threshold is based on the estimated head-start distance required for a sandpiper to accelerate from a standing start to a peregrine’s stealth attack velocity. Using measures of peregrine stealth attack velocity in Burns and Ydenberg (2002), we estimated the head-start distance using the method developed by Elliott et al. (1977) to calculate the distance within which lions had to approach wildebeests undetected in order to make a successful surprise attack. Hedenström and Rosén (2001) apply similar logic in models of prey escape strategies during falcon hunts (i.e. aerial climbing). We estimate the required head-start as 150 − 250m, at minimum. Most sandpiper foraging takes place further than this distance from shore (see Figure 1b in Pomeroy, 2006). Figure S2 reports the results of a sensitivity analysis of this assumption.

We calculated size and the safety indices from the CanVec map layers data set produced by Natural Resources Canada (acquired from:www.GeoGratis.gc.ca), which shows intertidal habitat and shoreline to a scale of 1:50,000. We extracted a polygon of intertidal as the waterbody features labelled as ‘temporary’ under the “Hydro” feature category within the CanVec dataset. We also extracted the highwater line layer and created a buffer of 150m around that line, which was then clipped to the intertidal layer. For each Universal Transverse Mercator (UTM) region, we transformed each polygon layer from original geographic projection (North American Datum (NAD) 1983 CSRS; Spheroid: GRS 1980; WKID: 4617) to the UTM region projection (UTM 19-22N WGS84) and clipped it to that grid. Around each site location we created a buffer 2500m in radius and defined the area of intertidal habitat as the area of the intertidal polygons that fell within that buffer (Figure 4).

### 2.4 Priority Matching Distribution index

We describe the distribution of sandpipers across locations in each year using a ‘Priority Matching Distribution’ (PMD) index. The PMD index assesses how closely the measured proportional distribution of sandpipers matches various distribution possibilities, ranging from sandpipers aggregating at dangerous locations, to spreading evenly over locations, to aggregating at safer locations (Ydenberg et al., 2017).

The PMD index is calculated as follows. In each year, *k* sites are surveyed and are indexed as *i* = 1, 2, 3, …, *k*, ordered from most dangerous (*i* = 1), to safest (*i* = *k*). The annual mean number (across all surveys) of sandpipers censused (‘usage’) at location *i* is denoted *U*_*i*_. The area of intertidal habitat at that location is denoted *A*_*i*_. The safety index for the site, *y*_*i*_, is the proportion of the site’s total intertidal area that lies more than 150m from the shoreline, whereas the danger index (*x*_*i*_ = 1 − *y*_*i*_) for the site is the proportion that lies within 150m of the shoreline (Figure 4).

We calculated the proportional area (*p*_*i*_) and bird usage (*q*_*i*_) of each location in relation to the total area surveyed and birds counted for all locations sampled in a given year. In each year, the cumulative proportion of the total area surveyed up to location *i* is

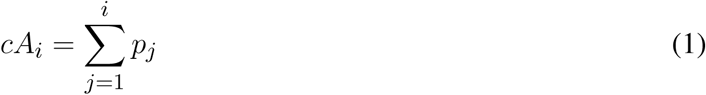

where the cumulative proportional area of all *k* sites surveyed in a year *cA*_*k*_ = 1. Analogously, the cumulative proportion of usage up to location *i* is calculated as

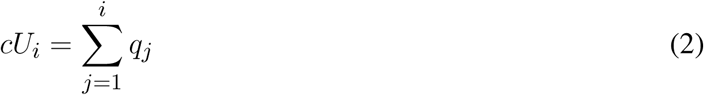

Calculation of the annual PMD index involves comparing the area under the curve (‘AUC’) of measured sandpiper usage (Equation 2), with that expected if sandpipers are distributed in relation to the intertidal area of each location (Equation 1; see Figure 5). AUC is calculated using a trapezoidal function. The trapezoid function for area of habitat surveyed is defined as

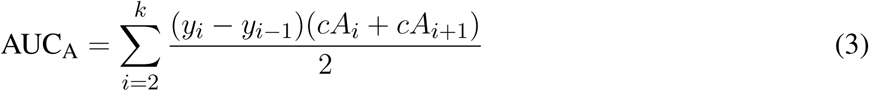

where *i* is a given location and *i* − 1 is the next most dangerous location. For bird usage the area under the distribution is calculated as

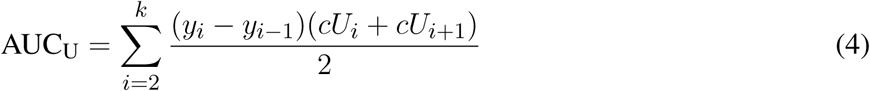

We used the trapezoid function because its estimate lies between that generated by the ‘upper-step’ and ‘lower-step’ functions. Sensitivity analyses using these step functions in place of the trapezoidal function produce only minor differences in the results.

The Priority Matching Distribution index is calculated as

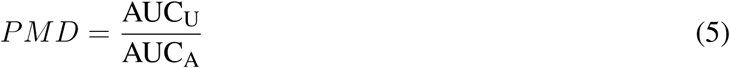

Values of the PMD index vary systematically with the distribution of sandpipers across locations, as summarized in Table 1 and shown in Figure S1. Note that the PMD index is calculated using proportions of the total number of sandpipers. It’s value is not affected by the number of sandpipers, unless the proportional distribution across locations also changes. Conversely, a change in the proportional distribution changes the PMD index value even if the total number of sandpipers remains unchanged.

**Table 1.** Simulated values of the PMD index as the usage distribution of semipalmated sandpipers over 100 simulated census locations. A bootstrap was used to estimate 95% CI intervals.

| Name | DESCRIPTION of DISTRIBUTION | lci | PMD index value | uci |
| --- | --- | --- | --- | --- |
| Danger Aggregation | 90% of usage at the most dangerous 20% of sites | 1.68 | 2.06 | 2.57 |
| Site Matching | Usage equal on all locations | 1.20 | 1.41 | 1.69 |
| Intermediate Aggregation | Aggregation on mid-safety locations | 0.93 | 1.29 | 1.78 |
| Area Matching | Usage proportional to intertidal area | 1.00 | 1.00 | 1.00 |
| Safe Area Matching | Usage proportional to safe intertidal area | 0.56 | 0.67 | 0.78 |
| Safety Aggregation | 90% of usage at the safest 20% of sites | 0.19 | 0.29 | 1.24 |

### 2.5 Analysis

Locations vary in the number of surveys, both within and between years (Figure 3). To examine the potential influence of individual locations we calculated the PMD index with and without each location (‘leverage’, see Table 2). Based on this, we excluded from the analysis 10 of 44 years that did not include surveys at one of the two most surveyed locations, namely Mary’s Point, NB (45.72°N, 64.65°W) and Johnsons Mills, NS (45.81°N, 64.5°W). We calculated and analyzed trends in the PMD with both locations excluded to ensure the results were not driven entirely by these locations (see Figure S2). We also excluded the year 1995, which had an extremely high count at a site surveyed in no other year that had a strong influence on the annual PMD. Our final data set included 3,030 surveys at 198 stopover locations, made 1974 − 2017 (excluding 1990, 1991, 1995, 1998, 2008, 2010, 2011, 2013, 2014).

**Table 2.** Sites with the most influence on the Priority Matching Distribution index across all years within the survey dataset. Leverage is defined here as the Cook’s distance of annual influence (sum of the squared differences between PMD calculated with and without a site, divided by the interannual variation in the PMD).

| Site Name | Safety ($y_i$) | Area ($km^2$) | Number of Years | Leverage |
| --- | --- | --- | --- | --- |
| Mary's Point | 0.84 | 7.90 | 27 | 426.03 |
| Grande Anse / Johnson's Mills | 0.88 | 6.95 | 18 | 232.54 |
| Saints Rest Marsh and Beach | 0.39 | 2.16 | 17 | 9.16 |
| Economy | 0.80 | 5.90 | 14 | 4.80 |
| Daniels Flats | 0.90 | 8.82 | 11 | 3.03 |
| Selma | 0.87 | 8.38 | 3 | 2.93 |
| Little Dyke | 0.87 | 6.71 | 4 | 2.68 |
| Cape Sable (Hawk Flats) | 0.59 | 3.36 | 28 | 2.61 |
| Egmont Bay (lower Bedeque area) | 0.81 | 4.00 | 1 | 2.54 |
| Daniel Head | 0.09 | 0.51 | 10 | 2.41 |
| Bedeque Bay | 0.73 | 5.07 | 7 | 2.37 |
| Cooks Beach | 0.38 | 6.03 | 28 | 1.58 |
| Lusbys Marsh/John Lusby Saltmarsh | 0.89 | 6.40 | 2 | 1.36 |

Our analysis focused on two questions: 1) how do semipalmated sandpipers distribute across stopover locations; and 2) has the proportional distribution of semipalmated sandpipers changed systematically since surveys began in 1974? We calculated the PMD index for each survey year. We examined annual change using a linear model, centred and rescaled by year to provide a meaningful intercept and provide a more accurate effect size (Gelman and Hill, 2006). We used Akaike Information Criterion (AIC) to compare support for a linear trend by competing a null model, a linear interannual trend model, a model with a quadratic term, and a model with the log of the interannual trend. We assessed the fit to the linear trend by bootstrapping the original count data to compute 95% confidence intervals of the intercept and interannual trend estimates.

We carried out a series of sensitivity analyses. To explore the sensitivity of the PMD index to our assumptions of location size (2.5 km) and the danger buffer (150 m), we altered the radius used to calculate *A*_*i*_ from 2.5km to 1km and 5km, and modified the danger buffer from 150m to 50m, 300m, and 450m. We also expanded the dates of surveys to include the 60th, 90th, 95th, 98th quantiles of dates, and with all surveys between July and October.

We recalculated the PMD in each year excluding Mary’s Point and Johnsons Mills. To control for the site bias towards a greater number of dangerous sites in later years (see Figure 6), we binned sites into 0.1 categories of safety. We sampled one site from each bin in each year, creating equal numbers of sites in all years. We resampled 1000 times and calculated the slope and intercept of the calculated PMD for each draw. We simulated the impact of sea level rise by reducing the total area of habitat available by the rate described in Murray et al. (2019), recalculating the safety index and redistributing total number of birds counted in that year across the site using a *Beta*(1, 14) distribution. For each simulation we calculated the PMD for each year and recalculated the intercept and rate of change in the linear interannual trend model.

**Figure 6.**
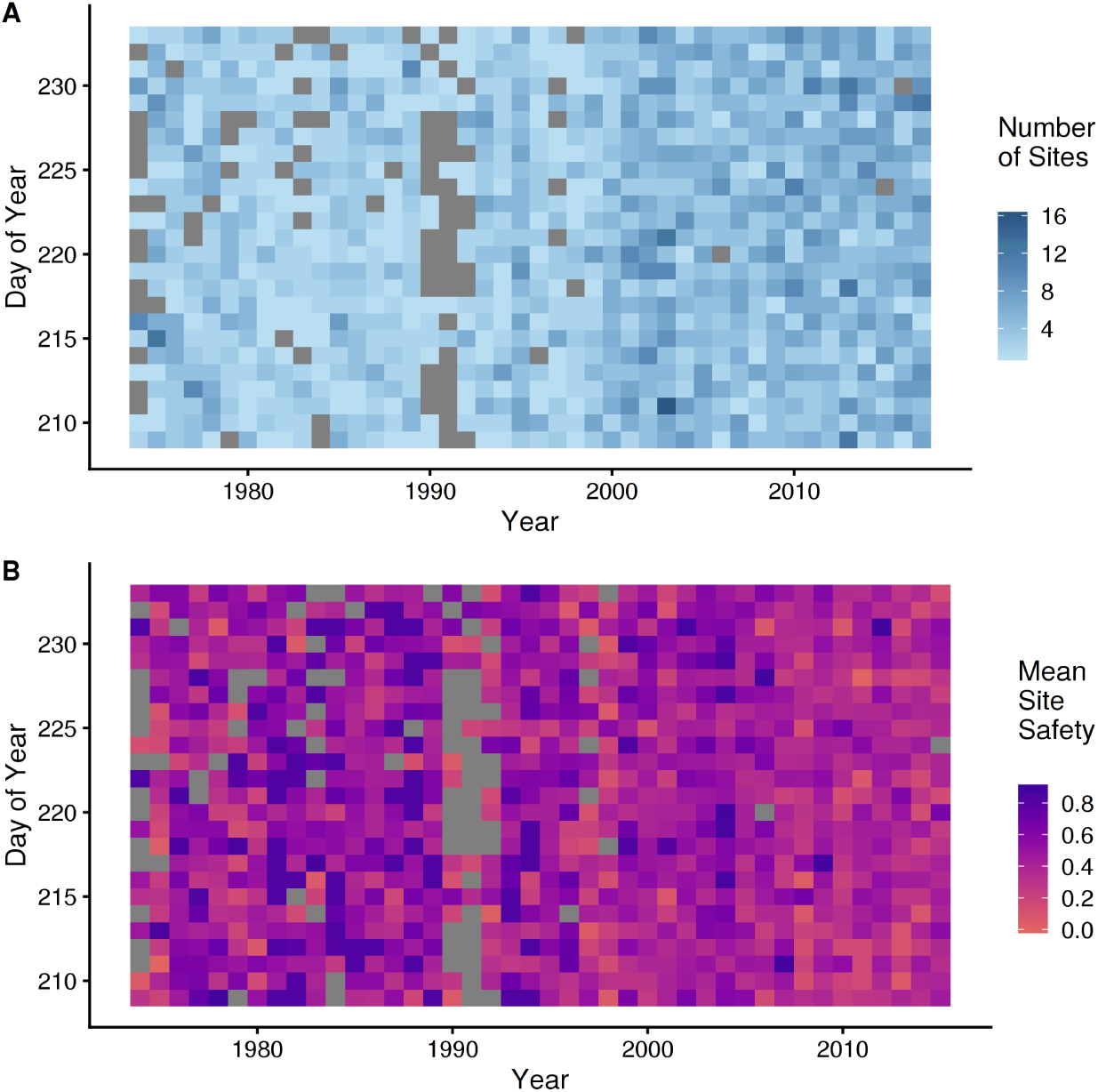
Biases and variation in site survey effort across years and dates. (A) Number of sites surveyed in a given day across years where darker blue shows more sites surveyed on a given day of year in a year. (B) Mean safety (*y*_*i*_) of sites surveyed in a given day across years where red indicates more dangerous sites surveyed for a given day and purple shows safer sites surveyed on that day. Grey represents zero surveys in both figures.

Finally, to assess whether the trend in the PMD index was driven by site size or site safety, we modified the calculation of the PMD index by arranging sites from smallest to largest instead of most dangerous to safest.

## 3 RESULTS

### 3.1 Location Characteristics

The 198 locations are arrayed over 4.6 degrees of latitude and 7.2 degrees of longitude (mean: 46°N, −64°W; Figure 2). On average, 27 locations were censused per year (range 12 − 44), averaging 3.2 surveys each per year (range 1.9-4.3). The total annual count (summed over all locations) of semipalmated sandpipers varies between 35,636 birds (1987) and 421,982 birds (1992) with a mean of 100,486, and no trend across years (*β*=0.0021 [−0.018, 0.021], using a log link).

Locations range in size from 0.002 km^2^ to 11 km^2^, with a mean of 3 km^2^. Safety indices ranged from 1.0 to 0.098 with a mean of 0.57. Most locations are small and relatively dangerous (Figure 7). There is overall a negative (log-linear) relation between location size and danger, so that large sites are on average safer, though note the wide variation. For example, locations 3 − 4 *km*^2^ in size have danger index values ranging from 0.2 − 0.8.

**Figure 7.**
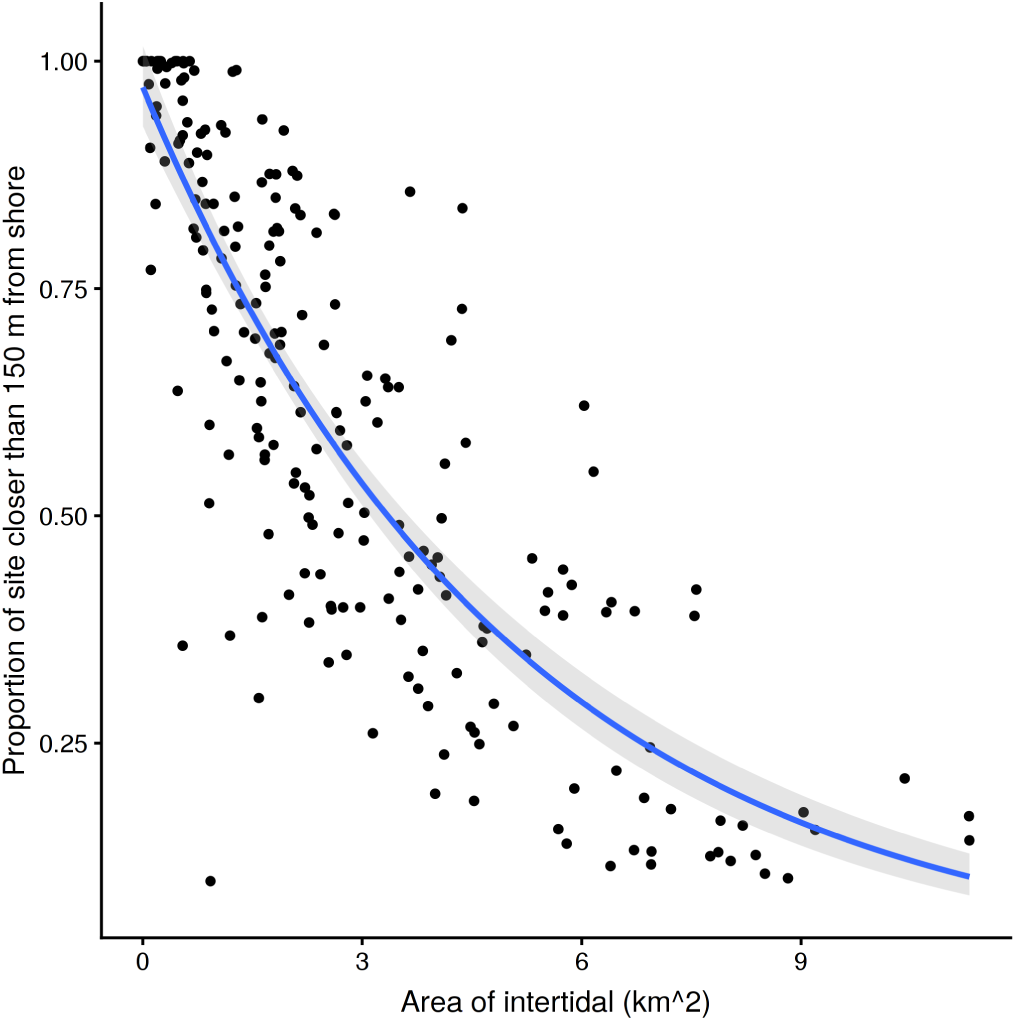
Location danger as a function of size. The fitted line shows the log-linear trend of danger with intertidal area. Larger sites are generally less dangerous, but the danger index varies widely between sites of a given size.

### 3.2 Sandpiper distribution

Annual PMD index values (Figure 8) range from 0.61 to 0.11 with an overall mean of 0.30 (95% CI [0.28, 0.36]). Most estimates of the PMD index are well below 0.50 indicating that semipalmated sandpipers aggregate at safer locations. The overall annual mean value of 0.30 corresponds with that expected when 90% of usage occurs at the safest 20% of locations (see Table 1).

**Figure 8.**
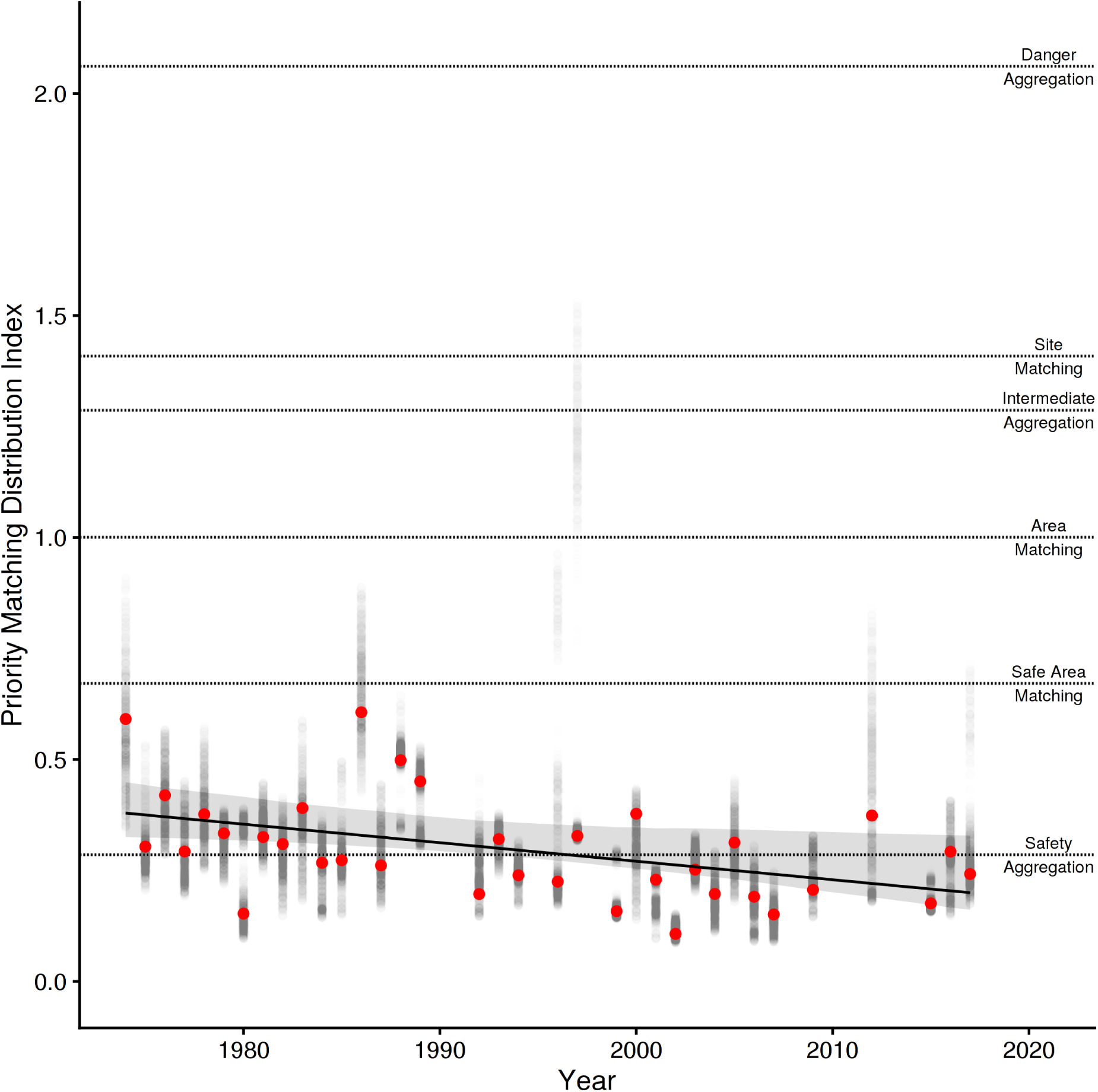
Interannual trend in the Priority Matching Distribution index (PMD) with the 95% bootstrapped confidence intervals. Points are the PMD for each year (red points) with the 95th inner quantiles of variation in the estimate shown in the grey points. The expected PMD index values for various distributions (danger aggregation, site-matching, area-matching, safe-area-matching, and safety aggregation) are shown as horizontally dashed lines.

Linear and log-linear models both estimate a decline in the PMD index over years (Table 3), and have approximately equal support from the data (*w*_*i*_ = 0.37 and *w*_*i*_ = 0.45 respectively). The quadratic model is less well supported (*w*_*i*_ = 0.16), and the null model not at all (*w*_*i*_ = 0.019). There is little deviation from linearity in the estimated trends for the log-linear model, and we therefore consider only the linear model. It shows that the PMD index falls at a standardized rate of −0.11 (95% CIs [−0.18, −0.034]) per SD of years (13 y), equivalent to −0.004 per year, for a 0.18-point decline in the PMD index between 1974 and 2017. This decline could be created either by (i) birds crowding into fewer sites (90% of birds at the 27% (1974) safest sites, shifting to the 13% (2017) safest sites); (ii) more birds crowding into the 20% safest sites (from 80% of birds in 1974 to 97% in 2017); (iii) or some combination of the two. Despite the extensive variability in methodology and the irregular coverage, the regression provides a reasonable fit (*r*^2^ = 0.21) to the data.

**Table 3.** Support for models of interannual trends in the Priority Matching Distribution. Models other than the null model use a centred and standardized variable for Year. The log-linear model is a linear generalized linear model with a log link function for the gaussian distribution. All other models are linear models with the identity link function. We used AICc values to correct for biases in the Akaike’s information criterion in models with low sample sizes, log(L) is the log-likelihood value, Δ_*i*_ is the difference in AICc value from that of the top model (i.e., lowest AICc), K is the number of parameters in each model and *w*_*i*_ is the Akaike weight. *r*^2^ are listed to show improvement of model fit between null and fitted model

| Interannual Trend | $\log(L)$ | $\Delta_i$ | K | $w_i$ | $r^2$ |
| --- | --- | --- | --- | --- | --- |
| Log Linear | 30.28 | 0.00 | 3.00 | 0.45 |  |
| Linear | 30.08 | 0.40 | 3.00 | 0.37 | 0.21 |
| Quadratic | 30.52 | 2.09 | 4.00 | 0.16 | 0.23 |
| Null | 25.94 | 6.29 | 2.00 | 0.02 | 0.00 |

AICc = -53.79; n = 35

The sensitivity analyses (Figure S2) demonstrate that variation in the number or danger of the sites surveyed each year does not bias the PMD index estimates, and is therefore unlikely to explain the interannual trend. Likewise, modification of the assumptions governing neither the selection of data nor those underlying the PMD calculation alter the results. ‘Binning’ the sites does not change the mean interannual trend of the PMD index (*β*=-0.07), but it does reduce the precision (95% CI[−0.16, 0.034]). This is not surprising, as we drew only one site per bin per year. Simulating a response to sea-level rise (i.e. sites becoming smaller) does not replicate the observed interannual trend in the PMD, indicating the level of observed sea-level rise observed across the years could not alone cause the shift between sites.

Most importantly, the “Area Sorted PMD” analysis reveals that the temporal trend is eliminated when locations are ranked ‘small to large’ rather than ‘dangerous to safe’ (interannual trend: 0.0049; 95% CI[−0.11, 0.12]). This shows that the decline in PMD is better explained by a shift to safer rather than to larger locations.

## 4 DISCUSSION

Our results show that the tendency of southbound semipalmated sandpipers to aggregate at the safest stopover locations has steadily increased since 1974. Sensitivity analyses establish that the shift appears to have been made specifically toward sites of higher safety, rather than to larger sites. The shift cannot be accounted for by inclusion (or exclusion) of the two sites exerting most leverage (Mary’s Point and Johnson’s Mills; see Table 2), by possible confounds arising from habitat reduction, by the selection of survey dates included in the analysis, or by altering our assumptions regarding the definitions of site size and danger (Figure S2).

We predicted this shift based on the well-established increase in continental falcon populations since the early 1970s. A similar redistribution was previously documented for wintering dunlins on North America’s Pacific coast (Ydenberg et al., 2017) and was also attributed to the large increase in falcon presence. Of particular note is the introduction of captive-reared peregrines in the 1970s and ‘80s to major stopover areas such as the Bay of Fundy (Dekker et al., 2011) and Delaware and Chesapeake Bays and their environs (Watts et al., 2015). With home ranges of 123 − 1175 km^2^ and a daily range of 23 km^2^ around breeding locations (Enderson and Craig, 1997; Jenkins and Benn, 1998; Ganusevich et al., 2004), the impact of breeding peregrines musty be widespread throughout both regions.

For most of the twentieth century, these regions were essentially predator-free during sandpiper passage, so stopover site choices and behaviour by migrant sandpipers could have been based primarily on food availability, with the danger posed by falcons ignored. Lank (1983) observed individual semipalmated sandpipers at Kent Island in the Bay of Fundy during the late 1970s so encumbered by fat that they were captured by gulls. Paralleling observations made on western sandpipers in the Strait of Georgia (Ydenberg et al., 2004), semipalmated sandpiper fuel loads in the Bay of Fundy have decreased at small, dangerous locations such as Kent Island, but not at large, safe locations such as Johnson’s Mills (Hope, 2010). The mass decline is attributed to the reduced predator escape performance induced by large fuel loads (Burns and Ydenberg, 2002), and is consistent with the hypothesis that stopover site choice and behaviour is strongly influenced by the trade off between fuel loading and predation danger (Pomeroy et al., 2008; Taylor et al., 2007).

Migrant sandpipers have previously been shown to be sensitive to predation danger on migration. The migratory behaviours responding to danger include flock size, vigilance, over-ocean flocking during high tides, length of stay at dangerous locations, location selection, habitat selection within a location, and fuel load (Dekker, 1998; Ydenberg et al., 2004; Pomeroy, 2006; Pomeroy et al., 2008; Sprague et al., 2008). Migrant sandpipers also change their behaviour seasonally in relation to their temporal proximity to the arrival of migrant peregrines (Hope et al., 2014, 2011). In a previous paper (Lank et al., 2017) we attributed the shortening wing length measured 1980 - 2015 in semipalmated sandpipers and other calidridines (Anderson et al., 2019) to selection for better predator escape performance (see also Ydenberg and Hope, 2019).

The PMD decline (Figure 8) has progressed steadily since 1974. The decline in PMD arose as the usage shown in Figure 5A shifted rightward, reducing the index value by 0.4% per year, for a total decline of 18% since 1974. It might be expected that the higher rate of increase in the number of breeding falcons in more recent years (Figure 1) should have accelerated the PMD decline. But the large majority of intertidal feeding area is on safer sites (see Figure 5B), with dangerous sites contributing disproportionately little to the total. We hypothesize that the initial small number of peregrines had a very large effect at small, dangerous locations (e.g. Page and Whitacre, 1975), where usage presumably began to drop in earlier years. The impact of additional peregrines was reduced as usage shifted to larger and safer sites.

Larger groups also have benefits in reducing the likelihood of being selected by a predator (dilution), and increased detection of predator attacks (many eyes Roberts, 1996; Bednekoff and Lima, 1998; Fernández-Juricic et al., 2007; Pays et al., 2013). With predation dilution can also come increased competition during foraging (Stillman et al., 1997; Vahl et al., 2005; Minderman et al., 2006). While the Bay of Fundy provides rich and widespread food for refuelling sandpipers, competitive interactions that reduce foraging efficiency likely occur at small scales (Vahl, 2005; Beauchamp, 2009, 2014). For most semipalmated sandpipers, the benefits of large aggregations appear to outweigh the costs to foraging efficiency.

Shorebird population census work indicates that many species, among them semipalmated sandpipers, have declined steadily since the 1970s (Bart et al., 2007; Andres et al., 2012; Gratto-Trevor et al., 2012; Smith et al., 2012; Morrison et al., 2012). Could the shift to safer stopover sites observed here be driven by this population decline? The PMD index is calculated based on proportions, so a reduction in sandpiper numbers would not affect the PMD value unless accompanied by a distributional change. Any distribution that includes safety considerations (Grand and Dill, 1999; Moody et al., 1996) would be expected to shift the towards safer sites as numbers decline even without an increase in predator numbers, due to the heightened danger of smaller numbers. The shift should progress until the fitness costs (reduced feeding rate) of the safer sites is compensated by the benefit (increased safety), which in turn depends on the marginal rates of change in food and safety with sandpiper density at each location (Ydenberg et al., 2017). Further evaluation is required.

Our analysis confirms the measured shift in the PMD index is better explained by a shift in distributions specifically toward safer and not just toward larger sites. The mean census numbers in the dataset used here show no temporal trend at all (see Results), but there has been a well-documented establishment of a large locally breeding population of peregrines. We are unable to exclude a possible contribution from population decline to the PMD change measured here, but all the evidence available is consistent with a strong non-lethal influence of predation danger.

An increase in food availability, or a reduction in energy demand would also allow greater aggregation on larger and safer sites, and thus a shift away from smaller and more dangerous locations. There is so far as we are aware no evidence for any trend in food abundance in the Bay of Fundy, nor is there any change proposed by the literature. The copepod *Corophium volutator*, is a major prey item for semipalmated sandpipers in the Bay of Fundy, and appears to vary in abundance between locations and within each year, but variation between years does not appear to be substantial (Barbeau et al., 2009). Other studies have shown variation between years when looking at a wider array of potential food sources (Quinn and Hamilton, 2012), but it appears that semipalmated sandpipers have flexibility in their food sources (Quinn et al., 2017), so that a decline at one location could be compensated by increases at others.

Other mechanisms could affect sandpiper energy requirements and thus affect distributions by reducing the need for food. A temperature increase due to climate change could reduce existence energy, though we note that the great majority of the intake of semipalmated sandpipers is used as fuel for the long trans-Atlantic flight to South America — which is not temperature dependent. Another possible climate change effect could operate by ecological mismatch (Jones and Cresswell, 2010). Southward migration timing is widely believed to match food at stopovers, but if the timing of the food peak has shifted due to climate change, the availability of food at stopovers could be affected. The result would be lower rather than higher food availability, which would, according to our current understanding, shift sandpipers to higher food (more dangerous, smaller) sites, opposite to the trend documented here.

In conclusion, semipalmated sandpipers aggregate in large numbers at a few large and safe sites and have steadily shifted towards safer sites between 1974 and 2017. The results appear robust to various biases possibly inherent in the dataset, and we suggest that the observed trend as a response to increased predator populations, especially the (re)introduction of predators at major stopover areas along the southbound migratory route. This result matches the previously reported shift in the non-breeding distribution of Pacific dunlins over the same period (Ydenberg et al., 2017), as well as the reduced stopover duration of southbound western sandpipers at dangerous stopover sites (Ydenberg et al., 2004). These adjustments likely have consequences for the schedule and routing of migration that we suspect may in turn contribute to substantial non-lethal effects on their populations.

## CONFLICT OF INTEREST STATEMENT

The authors declare that the research was conducted in the absence of any commercial or financial relationships that could be construed as a potential conflict of interest.

## AUTHOR CONTRIBUTIONS

DH, RY, and DL conceived of the study. PS and JP provided advice on the data use to fit with the study goals. JP organized data collection and cleaning. DH, with RY and DL designed the PMD index. DH performed the analysis, simulations. DH, RY, and DL wrote the paper with input from PS and JP.

## FUNDING

This study was supported by the Natural Sciences and Engineering Research Council of Canada, Environment and Climate Change Canada, and Simon Fraser University.

## ACKNOWLEDGEMENTS

The many of the members of the Centre for Wildlife Ecology at Simon Fraser University provided feedback on the development of the PMD index. NatureCounts and Bird Studies Canada managed and provided access to the ACSS dataset; we extend our sincere thanks to the many volunteers who contributed to this dataset. This paper was originally written as a thesis chapter and benefited from the feedback provided by DH’s committee members and thesis examining committee. A version of this manuscript was included in the PhD thesis of DH (Chapter 4 in Hope, 2018). This manuscript has been released as a Pre-Print at bioRxiv (Hope et al., 2019).

## Notes

#### Summary of Updates

Revised Introduction/Discussion. Added Figure 1 with Bay of Fundy Falcons

