## Supplementary Material for "Migrant semipalmated sandpipers (Calidris pusilla) have over four decades steadily shifted towards safer stopover locations"

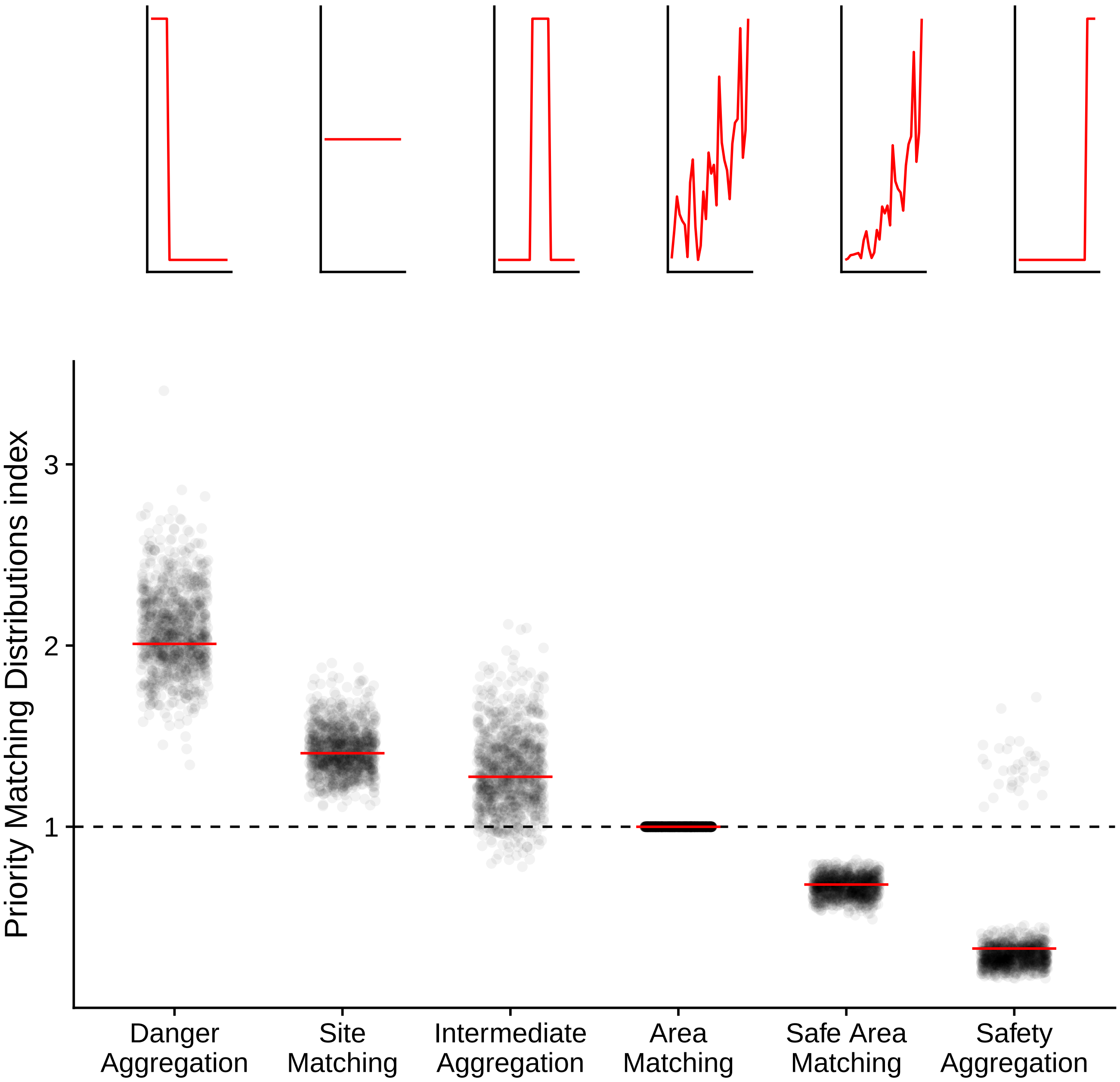


**Figure S1.** Simulations of birds across 30 sites using different distribution types. Distributions are described in text. Red line shows the baseline estimate of the PMD index for each distribution when site safety is sequentially assigned while the grey dots show the variation in the index when site safety is randomized. Points are jittered on the horizontal axis to aid in visualization. Above the points are examples of each distribution type showing sites arranged from dangerous to safe (left to right) and site abundance increasing vertically.

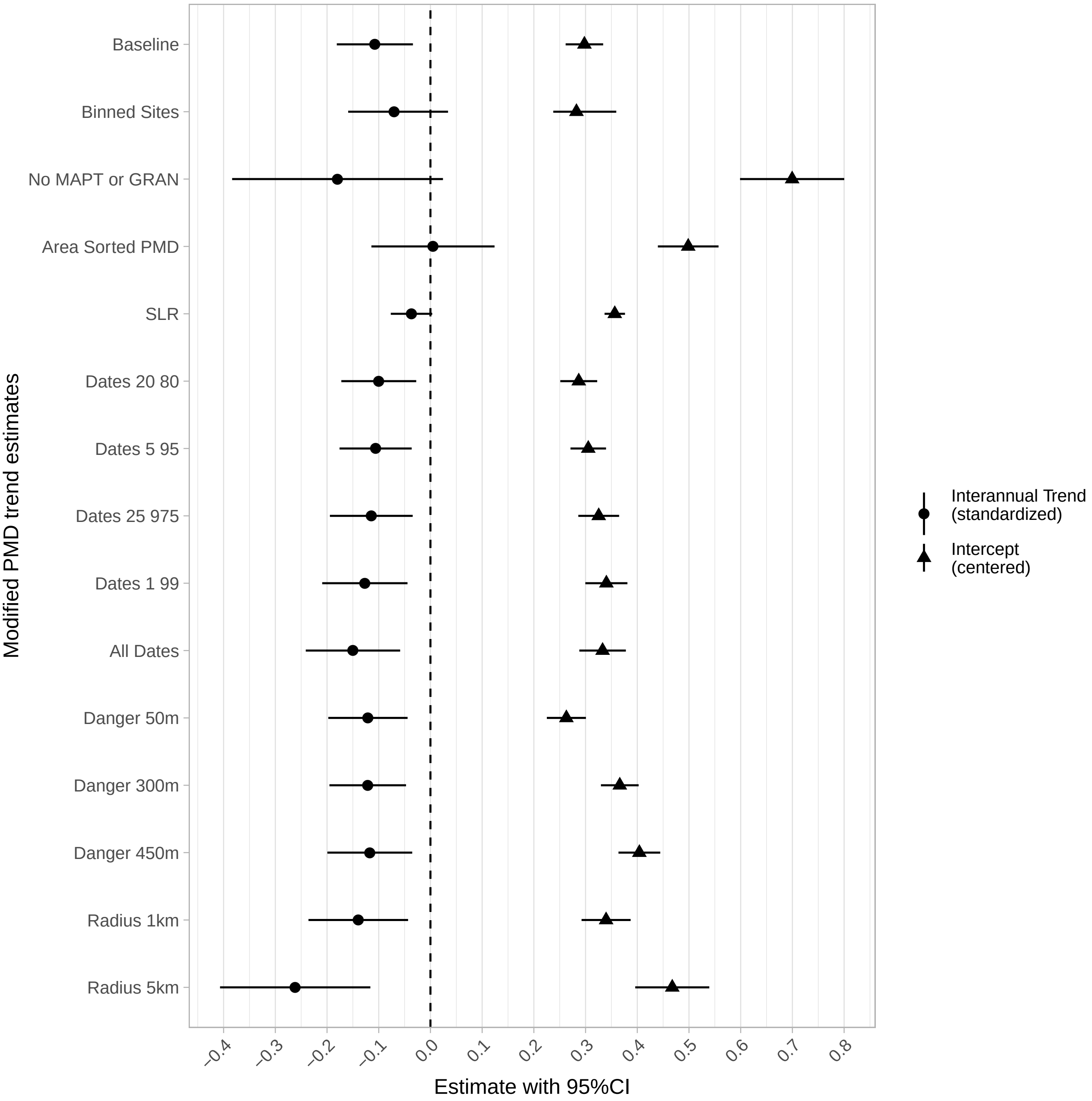


**Figure S2.** Intercept and standardized interannual trend estimates with accompanying 95% confidence intervals for repeated analyses where the assumptions underlying the priority matching distribution (PMD) index were relaxed. The "Baseline" run shows the main results shown from Figure 7. The "Binned Sites" analysis separated sites into bins of 0.1, sampled one site from each bin for each year and calculated interannual slopes and intercepts, repeating the process 1000 times. The "No MAPT or GRAN" analysis refitted the PMD trend model excluding the two sites exerting the most leverage (Mary's Point and Johnson's Mills; see Table 2). This did not affect the trend, but increased the intercept (i.e. less aggregation). The "Area Sorted PMD" run recalculated PMD by sorting by area of habitat rather than the safety index. Informatively, this eliminated the trend altogether, showing that the shift is toward greater safety rather than to larger size. The "SLR" simulation explored the result if birds were only responding to reductions in habitat. The "Dates" reanalyses clip the dates for inclusion in the analysis by the described percentiles. The "Danger" and "Radius" analyses recalculate the danger distance or the buffer around the geographic site location by the distance noted.

### Code availability

Code used in this analysis will be available at `https://github.com/dhope/pmd-sesa`. The packages used in this analysis are cited below.
